# Evidence of a common causal relationship between body mass index and inflammatory skin disease: a Mendelian Randomization study

**DOI:** 10.1101/265629

**Authors:** Ashley Budu-Aggrey, Ben Brumpton, Jess Tyrrell, Sarah Watkins, Ellen H Modalsli, Carlos Celis-Morales, Lyn D Ferguson, Gunnhild Åberge Vie, Tom Palmer, Lars G Fritsche, Mari Løset, Jonas Bille Nielsen, Wei Zhou, Lam C Tsoi, Andrew R Wood, Samuel E Jones, Robin Beaumont, Marit Saunes, Pål Richard Romundstad, Stefan Siebert, Iain B McInnes, James T Elder, George Davey Smith, Timothy M Frayling, Bjørn Olav Åsvold, Sara J Brown, Naveed Sattar, Lavinia Paternoster

## Abstract

**Objective:** Psoriasis and eczema are common inflammatory skin diseases that have been reported to be associated with obesity. However, causality has not yet been established. We aimed to investigate the possible causal relationship between body mass index (BMI) and psoriasis or eczema.

**Methods:** Following a review of published epidemiological evidence of the association between obesity and either psoriasis or eczema, Mendelian Randomization (MR) was used to test for a causal relationship between BMI and these inflammatory skin conditions. We used a genetic instrument comprising 97 single nucleotide polymorphisms (SNPs) associated with BMI. One-sample MR was conducted using individual-level data (401,508 individuals) from the UK Biobank and the Nord-Trøndelag Health Study (HUNT), Norway. Two-sample MR was performed with summary-level data (731,021 individuals) from published BMI, psoriasis and eczema GWAS. The one-sample and two-sample MR estimates were meta-analysed using a fixed effect model. To explore the reverse causal direction, MR analysis with genetic instruments comprising variants from recent genome-wide analyses for psoriasis and eczema were used to test if inflammatory skin disease has a causal effect on BMI.

**Results:** Published observational data show an association of greater BMI with both psoriasis and eczema case status. The observational associations were confirmed in UK Biobank and HUNT datasets. MR analyses provide evidence that higher BMI causally increases the odds of psoriasis (by 53% per 5 units higher BMI; OR= 1.09 (1.06 to 1.12) per 1 kg/m^2^; *P*=4.67×10^-9^) and eczema (by 8% per 5 units higher BMI; OR=1.02 (1.00 to 1.03) per 1 kg/m^2^; *P*=0.09). When investigating causality in the opposite direction, MR estimates provide little evidence for an effect of either psoriasis or eczema influencing BMI.

**Conclusion:** Our study, using genetic variants as instrumental variables for BMI, shows that higher BMI leads to a higher risk of inflammatory skin disease. The causal relationship was stronger for psoriasis than eczema. Therapies and life-style interventions aimed at controlling BMI or targeting the mechanisms linking obesity with skin inflammation may offer an opportunity for the prevention or treatment of these common skin diseases.

## INTRODUCTION

Psoriasis and eczema (atopic dermatitis) are two of the most common inflammatory skin disorders. Psoriasis affects approximately 2% of people within European populations[1] and is characterised by erythematous scaly plaques. Eczema affects 15-30% of children and 5-10% of adults in the developed world[2] and is characterised by itchy and inflamed skin. Obesity has become one of the leading health issues of the 21st century with over one quarter of the UK population now obese. In addition to clear links of obesity to diabetes and hypertension, observational evidence from epidemiological studies have suggested a relationship of increased weight with psoriasis[3] and more recently with eczema[4]. Furthermore, a small number of weight loss interventions have been shown to improve psoriasis and increase responsiveness to treatment[5-7]. Hypothetically, obesity could promote skin inflammation, but inflammatory skin disease could also lead to a reduction in activity levels, resulting in weight gain. A clearer understanding of the cutaneous and systemic metabolic effects associated with obesity and skin inflammation is an essential prerequisite to better define treatment and prevention strategies for these very prevalent public health issues.

Causality can be investigated with Mendelian Randomization (MR), which uses genetic variants to randomly allocate individuals to groups based on genotype (analogous to a randomised trial)[8]. Genetic variants are randomly allocated at conception, which proceeds disease onset. Therefore, confounding and reverse causation, common limitations of observational studies, can be avoided by using genetic variants as instrumental variables to estimate the causal effect of a risk factor upon an outcome of interest[8-10]. Genome-wide association studies (GWAS) have identified multiple genetic risk variants for complex human traits, including body mass index (BMI) for which 97 genetic risk loci have been identified to date, psoriasis (for which 63 loci have been identified) and eczema (24 loci within European populations)[11-13]. These studies have provided powerful data with which to perform MR[9].

In this study, we first reviewed the literature reporting observational evidence for associations between BMI and both psoriasis or eczema, and extended the observational associations in two large population based studies. We then applied MR to test for evidence of causality, the relevant strengths of associations, and to further define the direction of causality linking BMI with psoriasis or eczema.

## METHODS

### Literature Review

Studies were considered for review if they had reported on the association between BMI, obesity or being overweight and either psoriasis or eczema. We included studies with any operationalised definition of psoriasis or eczema, which could therefore include both atopic and non-atopic eczema as well as atopic dermatitis. For eczema, this work represents an update of the review from Zhang et al. from 2015[4]. PubMed was searched on 08/07/2016 with the terms “psoriasis AND (obesity OR overweight OR BMI)”, and on 18/11/2016 with the terms “(eczema OR atopic dermatitis) AND (obesity OR overweight OR BMI)”. Studies were included for review if they reported a statistical association between a BMI-related trait and either psoriasis or eczema. Results were extracted, and random effects meta-analyses were separately conducted for psoriasis and eczema, using the format of results presented by the majority of the papers reviewed. The meta-analyses were performed separately for children and adults, as well as for the groups combined.

For ease of comparison, where the combined result was the mean difference in BMI between cases and controls, this was transformed to an approximate expected effect of BMI on disease with the following formula, as previously demonstrated by Perry *et al*[14]:

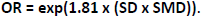

Where SD is the standard deviation increase in BMI per standard deviation change in the BMI genetic instrument (genetic risk score); SMD is the standardised mean difference and 1.81 is the scaling factor used to convert standardised mean differences to ln(ORs).

### Investigating causal relationships

#### Study populations

Data were available, with written informed consent for a total of 401,508 participants including 382,950 from the UK Biobank, aged between 40-69 years[15] (5,676 psoriasis cases; 9,933 eczema cases) and 18,558 from the Nord-Trøndelag Health Study (HUNT), aged 20+ years[16] (1,076 psoriasis cases; 472 eczema cases) (**Table 1**). All individuals were of European ancestry.

Summary level data were also available for 731,021 individuals of European ancestry from published GWAS studies for BMI[11] (n=322,154), psoriasis[12] (n=305,801) and eczema[13] (n=103,066).

**Table 1:** Descriptive statistics of datasets used in the study.

| Dataset | Sample size | Psoriasis cases/controls (prevalence) | Eczema cases/controls (prevalence) | Females (%) | Mean [SD] age (years) | Mean [SD] BMI (kg/m<sup>2</sup>) |
| --- | --- | --- | --- | --- | --- | --- |
| UK Biobank | 382,950 | 5,676 / 372,598 (1.5%) | 9,933 / 289,410 (3.3%) | 206,257 (54%) | 57 [8] | 27.4 [4.8] |
| HUNT | 18,558 | 1,076 / 17,145 (5.9%) | 472 / 16,951* (2.7%) | 10,076 (55.3%) | 53.7 [15.2] | 27.2 [4.4] |
| BMI GWAS<sup>[1]</sup> | 322,154 | - | - |  |  | 27.1 [4.6] |
| Eczema GWAS<sup>[2]</sup> | 103,066 | - | 18,900 / 84,166 (18.3%) |  |  |  |
| Psoriasis GWAS<sup>[3]</sup> | 305,801 | 21,399 / 95,464 (7.0%) | - |  |  |  |

BMI=body mass index; HUNT=Nord-Trøndelag Health Study; SD=standard deviation
\*Total sample size for eczema 17,423

#### Clinicai outcomes

##### - UK Biobank

The BMI of UK Biobank participants was calculated from standing height and weight measures that were taken while visiting an assessment centre. Units of measurement are kg/m^2^. Individuals were defined as having psoriasis or eczema based on their response during a verbal interview with a trained member of staff at the assessment centre. Participants were asked to tell the interviewer which serious illnesses or disabilities they had been diagnosed with by a doctor, and were defined as psoriasis or eczema sufferers if either disease was mentioned. Disease information was also obtained from the Hospital Episode Statistics (HES) data extract service where health-related outcomes had been defined by International Classification of Diseases (ICD)-10 codes (**Supplementary Table 1**).

Additionally, if any had answered “yes” to “Has a doctor ever told you that you have hay fever, allergic rhinitis or eczema”, then these individuals were excluded from eczema controls.

##### - HUNT

Participants' height and weight were measured and used to calculate BMI (kg/m^2^). Participants were defined as psoriasis or eczema cases based on their response to a general questionnaire sent to all HUNT participants. Psoriasis cases responded affirmatively to the question “Have you had or do you have psoriasis?”. The diagnostic properties of the psoriasis question have been validated in HUNT (positive predictive value was 78%; 95% CI 69 to 85)[17]. Eczema cases responded affirmatively to both “Have you had or do you have any of the following diseases: Eczema on hands” and “Did you have eczema when you were a child? (also called atopic eczema)”.

#### Genotyping

##### - UK Biobank

Genotyping of UK Biobank participants was performed with one of two arrays (The Applied Biosystems™ UK BiLEVE Axiom™ Array (Affymetrix) and Applied Biosystems™ UK Biobank Axiom™ Array). Sample quality control (QC) measures included removing individuals who were duplicated and highly related (3rd degree or closer), had sex mismatches, as well as those identified to be outliers of heterozygosity and of non-European descent. Further details of the QC measures applied and imputation performed have been described previously[18-21].

##### - HUNT

Genotyping of the HUNT participants was performed with one of three different Illumina HumanCoreExome arrays (HumanCoreExome12 v1.0, HumanCoreExome12 v1.1 and UM HUNT Biobank v1.0). Sample QC measures were similar to those applied to the UK Biobank. Related individuals were excluded (n=30,256), resulting in 18,558 individuals for analysis. Details of the genotyping, QC measures applied and imputation have been described elsewhere[22].

#### Statistical analyses

##### Observational analysis

Within the UK Biobank and HUNT datasets, logistic regression models were used to estimate the observational association between BMI and psoriasis, and BMI and eczema. Analyses were adjusted for age and sex and the estimates for each dataset were meta-analysed assuming a fixed effect model.

#### Genetic instruments

##### - BMI

The genetic instrument for BMI comprised the 97 BMI associated SNPs reported by the GIANT consortium (a meta-analysis of 125 GWAS studies with 339,224 individuals)[11]. These SNPs were extracted from both the UK Biobank and HUNT datasets.

We also combined these SNPs to create a standardised genetic risk score (GRS) using the ‑‑score command in PLINK (version 1.9). In doing so the dosage of the effect allele for each SNP was weighted by the effect estimates reported for the European sex-combined analysis (n= 322,154) by Locke *et al*[11], summed across all variants, and divided by the total number of variants in the calculation. The scores were standardized to have a mean of 0 and standard deviation of 1.

The BMI-associated SNP rs12016871 was not present within the UK Biobank and HUNT datasets, therefore rs9581854 was used as a highly correlated proxy (r^2^ = 1.0) (**Supplementary Table 2**).

##### - Psoriasis

62 psoriasis associated SNPs (outside of the human leukocyte antigen (HLA) region) were obtained from the most recent psoriasis GWAS study (a meta-analysis of 19,032 cases and 286,769 controls of European ancestry)[12]. These SNPs were extracted from both the UK Biobank and HUNT datasets, and combined to create a standardised GRS, where they were weighted by their published effect sizes. The psoriasis-associated SNP rs118086960 was not present in the UK Biobank imputed dataset and had no suitable proxy (r^2^ > 0.8). Therefore 61 independent SNP associations were used as a genetic instrument in this dataset (**Supplementary Table 2**).

##### - Eczema

We selected 24 SNPs that had been reported to be associated with eczema in white European populations in the most recent GWAS (meta-analysis of 21,399 cases and 95,464 controls)[13]. These SNPs were extracted from the UK Biobank and HUNT datasets and combined to create a standardised GRS, where they were weighted by the published effect estimates reported for European individuals (18,900 cases; 84,166 controls)[13].

Within the HUNT dataset, the eczema-associated SNP rs12153855 was not present and no suitable proxy (r^2^ > 0.8) was found, therefore 23 SNPs were used to create the genetic instrument in this dataset.

#### Mendelian Randomization analysis

One-sample MR analysis was performed separately in the UK Biobank and HUNT datasets with the individuals BMI SNPs, measured BMI and disease outcome status (**Figure 1a**). The MR estimates from each genetic instrument (SNP) were meta-analysed assuming a random effects model, giving a single estimate for the analysis performed in each dataset. This analysis was performed with the two-stage predictor substitution (TSPS) method[23]. The first stage involved regression of BMI upon individual BMI SNPs. The outcome (psoriasis or eczema) was then regressed upon the fitted values from the first regression stage. As eczema and psoriasis are binary outcomes, the first stage linear regression was restricted to individuals that were controls for the outcome only. Logistic regression was then performed in the second stage where the fitted values for the cases were predicted. The standard errors (SE) of these estimates were adjusted using the first term of the delta method expansion for the variance of a ratio, allowing for the uncertainty in the first regression stage to be taken into account[24].

**Figure 1:**
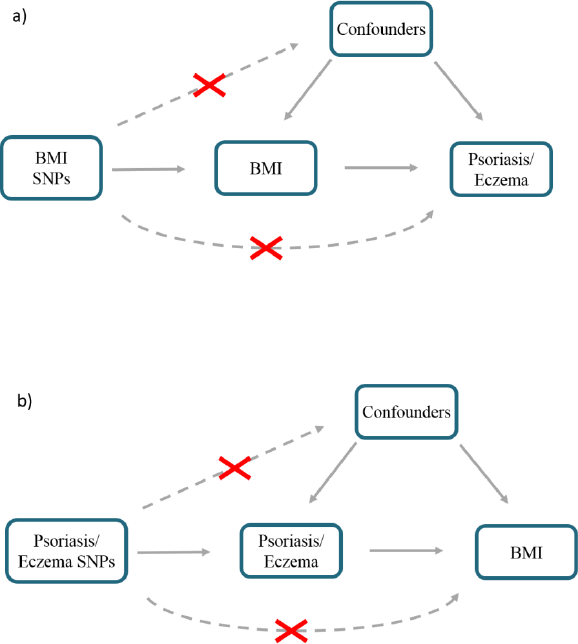
Schematic representation of MR analyses. a) BMI SNPs were used as instrumental variables to investigate the causal effect of BMI upon eczema and psoriasis. b) Eczema SNPs and psoriasis SNPs were used as instrumental variables to investigate the causal effect of genetic risk of psoriasis and eczema upon BMI. Crosses indicate violations of MR assumptions that there is no association between the instrumental variable and the confounders; and there is no presence of pleiotropy. BMI= body mass index; SNP=single nucleotide polymorphism.

Genetic principal components (as previously described [20-22]) were included as covariates in the analysis to control for residual population structure. UK Biobank analysis also controlled for the platform used to genotype the samples. In the HUNT dataset the genotyped data was harmonised before imputation.

Two-sample MR analysis of published GWAS data was performed using the “MendelianRandomisation” R package[25,26]. Estimates for the association between BMI and BMI SNPs in Europeans were taken from the GIANT BMI GWAS study[11]. Summary statistics from the most recent psoriasis[12] and eczema[13] GWAS studies were used to obtain estimates for the association of eczema and psoriasis with the BMI SNPs in Europeans. The published BMI SNP estimates were based on an inverse normal transformation of BMI residuals on age and age^2^, as well as any necessary study-specific covariates. In unrelated individuals, residuals were calculated according to sex and case/control status, and were sex-adjusted amongst related individuals[11]. Therefore the causal estimates for the two-sample analysis were converted to raw BMI units (kg/m^2^), assuming a median BMI standard deviation of 4.6 kg/m^2^[11].

The one and two-sample estimates were meta-analysed assuming a fixed effect model to obtain overall causal estimates.

#### Sensitivity analysis

MR-Egger regression, weighted median analysis and the weighted mode-based estimate (MBE) were used to investigate potential pleiotropy. SNPs that act through a pleiotropic pathway would violate the MR assumption that the instrumental variable has an effect upon the outcome only via the exposure being investigated, and could bias the causal estimate (**Figure 1**). The weighted median method provides a valid causal estimate if at least 50% of the information each instrument contributes to the analysis comes from valid instruments[27]. Likewise, the weighted MBE also provides a valid causal estimate if the largest weights are from valid instruments[28], whilst the intercept from the MR-Egger regression analysis allows the size of any pleiotropic effect to be determined[29].

In addition, one-sample MR analysis was performed using the *FTO* SNP alone (rs1558902) as a genetic instrument due to its strong association with BMI[30].

As the instrumental variables used in an MR analysis are assumed to be independent of confounders, we investigated the relationship between the BMI GRS and potential confounders of BMI by performing a simple regression of the confounder upon the BMI GRS. The relationship between the *FTO* variant and potential confounders was also investigated.

#### Reverse direction MR analysis

We investigated if either psoriasis or eczema has a causal effect upon BMI (**Figure 1b**). One-sample MR analysis was performed in the UK Biobank and HUNT datasets using the two-staged least squares (TSLS) method with the “ivpack” R package[31] using individual SNPs for each exposure as an instrument. This analysis involves two linear regression stages where psoriasis or eczema are first regressed upon the instrument (disease-associated SNPs), then the outcome (BMI) is regressed upon the fitted values from the first stage regression. The estimates from each genetic instrument were meta-analysed assuming a random effects model to give a single estimate for the effect of psoriasis and of eczema. Two-sample MR analysis was also performed, using summary results from GWAS studies for psoriasis and eczema[12,13] and from the GIANT BMI GWAS[11]. The one and two-sample MR estimates were meta-analysed using a fixed effect model.

Sensitivity analyses were performed with MR-Egger regression, weighted median and weighted MBE methods.

All analyses were performed using R (www.r-project.org) unless otherwise stated.

## RESULTS

### Literature review

#### Observational association of BMI and psoriasis

We identified 53 studies reporting on the relationship between BMI, obesity or being overweight and psoriasis. Of these, 34 compared mean BMI between psoriasis cases and controls (**Figure 2**). A meta-analysis of these found a mean difference in BMI between psoriasis cases and controls to be 1.26 kg/m^2^ (95% CI 1.01 to 1.51) amongst adults (69,704 psoriasis cases and 617,704 controls) and 1.55 kg/m^2^ (95% CI 1.13 to 1.98) in children (844 psoriasis cases and 709 controls). For both adults and children, the observed difference in BMI is equivalent to a 9% increase in the odds of psoriasis per 1 kg/m^2^ increase in BMI. 19 other studies tested for an association between BMI or obesity traits and psoriasis using alternative models[32-50]. These all reported a positive association, including two studies which reported the odds of psoriasis in adults per 1 kg/m^2^ increase in BMI to be 1.09 (95% CI 1.04 to 1.16)[51] and 1.04 (95% CI 1.02 to 1.10)[52].

**Figure 2:**
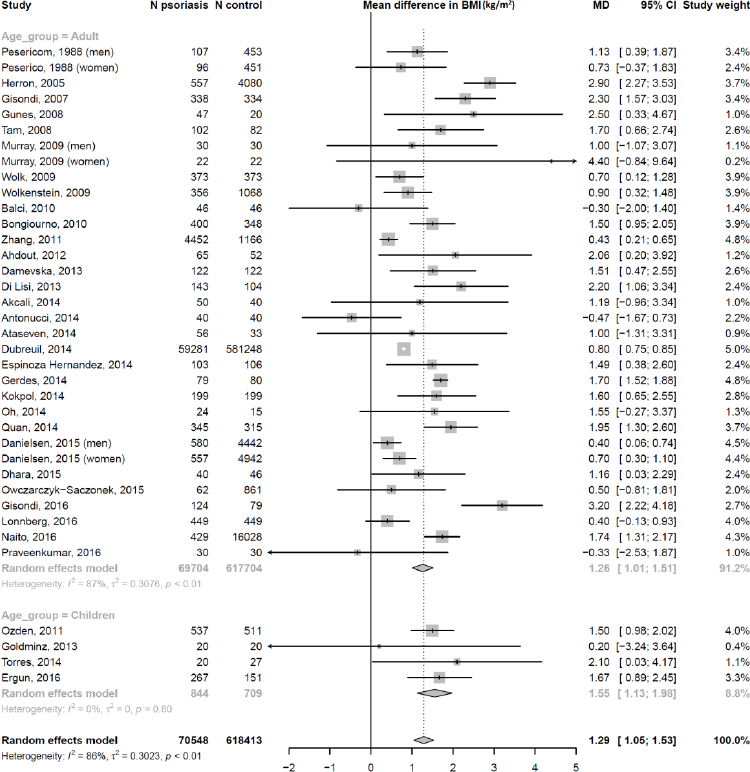
Observational association between BMI and psoriasis: meta-analysis of mean difference in BMI (kg/m^2^) between psoriasis cases and controls. BMI=body mass index; CI=confidence interval; MD=mean difference.

#### Observational association of BMI and eczema

We identified 42 studies that had investigated the relationship between BMI, obesity and being overweight and eczema. 5 studies reported the mean difference in BMI between eczema cases and controls[53-57]. These studies showed mixed results, with the majority reporting mean BMI to be lower in eczema cases compared to controls. Kusunoki *et al* reported the odds of eczema per 1 unit increase in BMI (1.02 per 1 kg/m^2^ increase in BMI)[54]. As the majority of studies (37 out of 42) report the OR for eczema in obese and/or overweight individuals compared to normal weight individuals, we updated the Zhang *et al*[4] meta-analysis and included these additional results. The definitions for obesity and being overweight were based on international guidelines[58] or population specific thresholds. For overweight individuals, the overall OR was 1.05 (95% CI 0.94 to 1.19) in adults (n=51,008) and 1.08 (95% CI 1.00 to 1.16) in children (n=506,202), compared to normal weight individuals (**Figure 3a**). For obese individuals, the corresponding OR of having eczema was 1.19 (95% CI 0.95 to 1.49) in adults (n=1,400,679) and 1.20 (95% CI 1.11 to 1.30) in children (n=796,514) (**Figure 3b**).

**Figure 3a:**
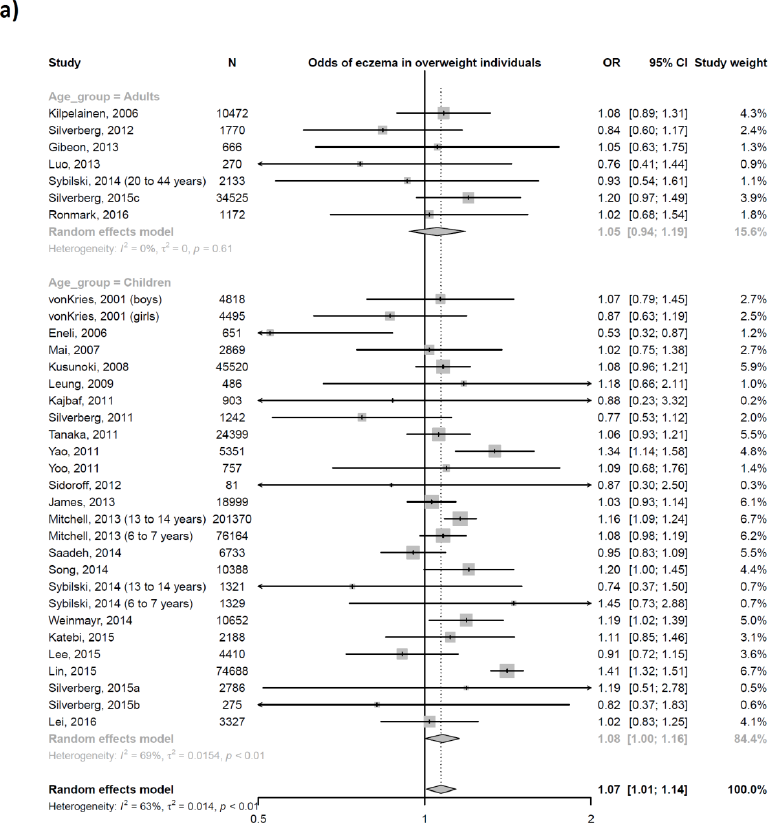
Observational association between increased weight and eczema: meta-analysis of the odds of eczema in overweight individuals compared with normal weight individuals. CI=confidence interval; OR=odds ratio.

**Figure 3b:**
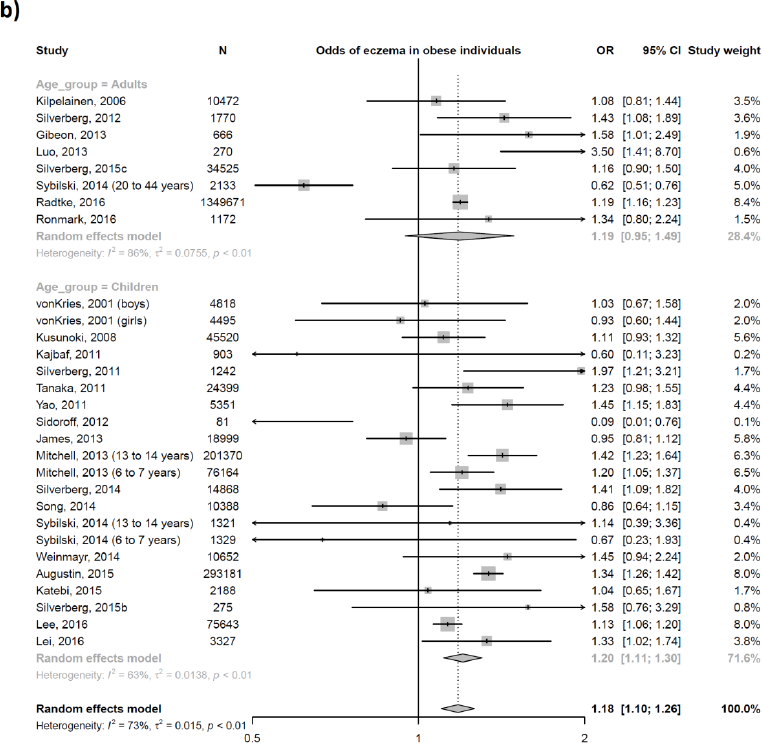
Observational association between increased weight and eczema: meta-analysis of the odds of eczema in obese individuals compared with normal weight individuals. CI=confidence interval; OR=odds ratio.

### BMI instrument

The BMI GRS was strongly associated with BMI in UK Biobank (Beta=0.64; 95% CI 0.63 to 0.66, F-statistic=7091, R^2^=1.8%) and HUNT (Beta=0.65; 95% CI 0.59 to 0.71, F-statistic=90.82, R^2^=3.4%), providing evidence in support of this instrument (**Supplementary Figures 1 and 2**). We investigated the association between the BMI GRS and potential confounders of BMI. Some small effects on the confounders were seen, however the strength of association was minimal in comparison to the association with BMI. This was also true for the *FTO* variant, which was found to have a much stronger association with BMI in UK Biobank (Beta=0.36; 95% CI 0.34 to 0.38, F-statistic=1152.1, R^2^=1.7%) and HUNT datasets (Beta=0.35; 95% CI 0.26 to 0.44, F-statistic=38.3, R^2^=1.4) compared to that with the potential confounders (**Supplementary Figures 3 and 4**).

### Effect of BMI upon psoriasis

#### Observational analysis

Higher BMI was associated with increased risk of psoriasis in both the UK Biobank and HUNT datasets. Overall, 1 kg/m^2^ increase in BMI was associated with 4% higher odds of psoriasis (metaanalysis OR=1.04; 95% CI 1.04 to 1.05; P=7.09×10^-68^) (**Figure 4**).

**Figure 4:**
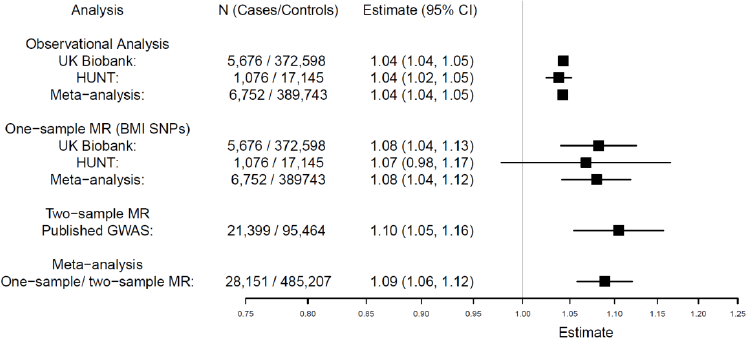
Effect of BMI upon psoriasis. Meta-analysis of one-sample and two-sample MR estimates using individual BMI SNPs as instrumental variables. Estimates are given per 1 kg/m^2^ increase in BMI. CI=confidence interval; MR=Mendelian Randomization; OR=odds ratio.

#### Mendelian Randomization

MR performed with UK Biobank, HUNT and published GWAS data gave evidence that higher BMI increases the risk of psoriasis. The causal estimate from UK Biobank showed an ∼8% increase in odds of psoriasis per 1 kg/m^2^ higher BMI (OR=1.08; 95% CI 1.04 to 1.13; *P*=8.75×10^-5^), while a ∼7% increase was shown in HUNT (OR=1.07; 95% CI 0.98 to 1.17; *P*=0.14). The two-sample estimate from published GWAS data also gave evidence of higher psoriasis risk with increased BMI (OR=1.10, 95% CI 1.05 to 1.16; *P*=6.46×10^-5^) (**Figure 4**). Meta-analysis of both one-sample and two-sample estimates produced an overall causal estimate of 1.09 per 1 kg/m^2^ higher BMI (95% CI 1.06 to 1.12; *P*=4.67×10^-9^) (**Figure 4**). There was little evidence of pleiotropy in the MR-Egger regression analysis (UK Biobank intercept=0.00; 95% CI -0.01 to 0.01; *P*=0.63, HUNT intercept=0.00; 95% CI -0.02 to 0.02; *P*=0.96) and the sensitivity analyses gave similar results (**Supplementary Figure 5, Supplementary Table 3**). In addition, when limiting the instrument to only the *FTO* SNP, a similar result (although with a wider confidence interval) was observed (OR=1.11; 95% CI 1.04 to 1.19; *P*=1.22×10^-3^) (**Supplementary Figure 6**).

### Effect of BMI upon eczema

#### Observational analysis

Observationally there was very little evidence of a relationship between BMI and eczema in the UK Biobank or HUNT datasets (**Figure 5**) (meta-analysis OR=1.00; 95% CI 0.99 to 1.00; *P*=0.31).

**Figure 5:**
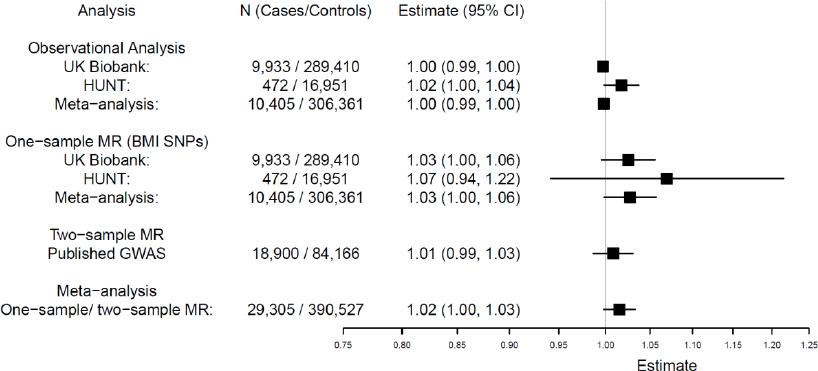
Effect of BMI upon eczema. Meta-analysis of one-sample and two-sample MR estimates using individual BMI SNPs as instrumental variables. Estimates are given per 1 unit increase in BMI (kg/m^2^). CI=confidence interval; MR=Mendelian Randomization; OR=odds ratio.

#### Mendelian Randomization

The causal estimates in UK Biobank (OR=1.03; 95% CI 1.00 to 1.06; *P*=0.10) and HUNT (OR=1.07; 95% CI 0.94 to 1.22; *P*=0.30) were higher than those seen observationally, suggesting an increased risk of eczema with higher BMI (OR per 1 kg/m^2^= 1.03; 95% CI 1.00 to 1.06; *P*=0.07). The two-sample estimate with published GWAS data was lower (OR=1.01; 95% CI 0.99 to 1.03; *P*=0.47). However, the meta-analysis of the UK Biobank, HUNT and two-sample estimates showed evidence of a small causal effect, increasing the odds of eczema by ∼2% (OR=1.02; 95% CI 1.00 to 1.03; *P*=0.09) (**Figure 5**). There was little evidence of pleiotropy in the MR-Egger regression analysis (UK Biobank intercept= 0.00; 95% CI -0.01 to 0.01; *P*=0.72, HUNT intercept=-0.01; 95% CI -0.03 to 0.02; *P*=0.71) and the sensitivity analyses gave similar results (**Supplementary Figure 7, Supplementary Table 3**). MR analysis with the *FTO* SNP alone gave a slightly stronger estimate, although with a wider confidence interval (OR=1.05; 95% CI 1.00 to 1.11; *P*=0.04) (**Supplementary Figure 6**).

### Reverse MR analysis

#### Effect of psoriasis upon BMI

The one-sample MR estimates in UK Biobank and HUNT gave little evidence that genetic risk of psoriasis has a causal effect on BMI, representing a 0.27 kg/m^2^ causal difference in BMI between cases and controls and a wide confidence interval (95% CI -2.02 to 2.55; *P*=0.82) (**Figure 6**). When combined with the two-sample estimate obtained using published GWAS data, there was weak evidence of a small causal effect of psoriasis on BMI (0.05 kg/m^2^ higher BMI in cases compared with controls, 95% CI 0.01 to 0.10; *P*=0.01), but this was not clinically relevant. We note that when performing the two-sample analysis, BMI summary statistics were only available for 55 of the 62 psoriasis SNPs.

**Figure 6:**
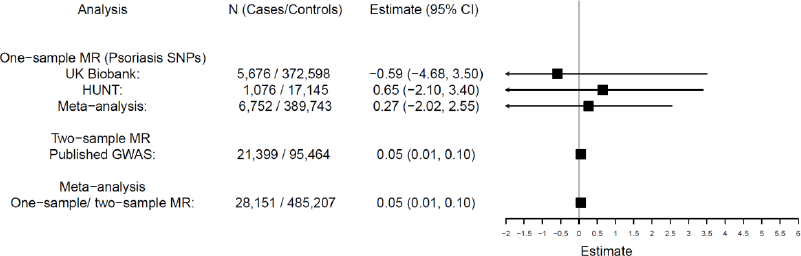
Reverse direction MR analysis - effect of psoriasis upon BMI: Meta-analysis of one-sample and two-sample MR estimates using individual psoriasis SNPs as instrumental variables. Estimates represent mean difference in BMI (kg/m^2^) between psoriasis cases and controls. CI=confidence interval; MR=Mendelian Randomization. F-statistic = 1316, R2 = 2.6% (UK Biobank); F-statistic = 394, R^2^ = 2.1% (HUNT).

#### Effect of eczema upon BMI

The one-sample MR estimates in UK Biobank and HUNT gave little evidence that genetic risk of eczema influences BMI, representing a 0.00 kg/m^2^ causal difference in BMI between cases and controls (95% CI -2.39 to 2.38; *P*=1.00) (**Figure 7**). Combined with the two-sample estimate obtained using published GWAS data there was some evidence that eczema could cause a very weak increase in BMI (0.10 kg/m^2^; 95% CI -0.01 to 0.20; *P*=0.07) (**Figure 7**). We note that the *FLG* loss-of-function variant (R501X/rs61816761) known to be strongly associated with atopic eczema[59] was not available in the two-sample analysis.

**Figure 7:**
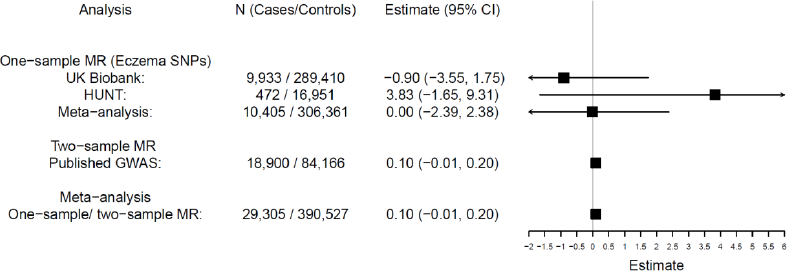
Reverse direction MR analysis - effect of eczema upon BMI: Meta-analysis of one-sample and two-sample MR estimates using individual eczema SNPs as instrumental variables. Estimates represent mean difference in BMI (kg/m^2^) between eczema cases and controls. CI=confidence interval; MR=Mendelian Randomization. F-statistic = 4722, R^2^ = 1.6% (UK Biobank); F-statistic = 1225, R^2^ = 6.5% (HUNT).

There was little evidence of pleiotropy in the causal estimates for psoriasis or eczema on BMI (**Supplementary Table 4, Supplementary Figures 8 and 9**).

## DISCUSSION

The rising prevalence of psoriasis[60], atopic eczema[61] and obesity[62] are important public health concerns. We found evidence of increased BMI having a clinically relevant causal effect upon psoriasis. The OR of 1.09 for one unit higher BMI suggests, for example, that an increase in BMI of 5 units would increase the risk of psoriasis by 53% (OR per 5 units higher BMI = exp(Beta per 1 unit higher BMI * 5). Furthermore, the direction and magnitude of effect seen for psoriasis is consistent with that seen observationally and in previous literature. In the reverse direction, the estimate of 0.05 kg/m^2^ demonstrates that there is little meaningful influence of psoriasis risk on an individual’s BMI, suggesting the observed association is predominantly explained by a causal effect of BMI upon psoriasis. Our analyses found little evidence for an observational association between BMI and eczema in this population. This is consistent with the summary of previous literature that showed strong evidence of an observational relationship between eczema and increased BMI in paediatric but not in adult studies. However, the MR analysis showed a small but potentially important effect of BMI upon eczema. The OR of 1.02 suggests that an increase in BMI of 5 units would increase eczema risk by approximately 8%. However, as shown with psoriasis, there was little evidence for a clinically meaningful effect of eczema on BMI (0.10 kg/m^2^).

We performed various sensitivity analyses to explore potential pleiotropic effects of the SNPs that make up the BMI instrument. When restricting the instrument to only the *FTO* variant, for which there is good understanding of the biological mechanism[30], we found the estimate from this analysis to be consistent with the estimate using all BMI SNPs. This suggests that the causal estimates seen are not driven by pleiotropic SNPs with unknown biological effects. This is supported by the MR-Egger regression intercepts which were centred around zero (indicating no directional pleiotropy amongst the included variants).

Our analysis has included a total of 1,132,529 individuals, including data from two of the biggest population based studies, and one of the largest published GWAS studies. We have applied both a one-sample and two-sample MR approach and the estimates from these analyses were meta-analysed to provide increased statistical power. The use of a strong genetic instrument for BMI provides an additional strength to this study. However, as expected with a large sample size, we did observe some associations between the BMI GRS and potential confounders of BMI. Nonetheless, we found these to be minimal in comparison to the strength of the association with raw BMI and therefore unlikely to be materially affecting the results. It is still important to note the possible influences of unmeasured confounders, especially when utilising large data resources such as the UK Biobank. The data in our study included contemporaneous measurements of BMI but relied predominantly on patient report for the ascertainment of psoriasis and eczema. Both of these diseases may follow a chronic relapsing-remitting or resolving course, but approximately 60% of atopic eczema shows remission in childhood[63] and this phenotype may therefore be more susceptible to recall bias in adult studies. This may have reduced the power to detect a causal association between BMI and eczema. In addition, the BMI SNPs used are a stronger instrument for adult BMI compared to childhood BMI[64]. Furthermore, the possibility of misclassification of inflammatory skin disease (psoriasis reported as eczema, or *vice versa)* and mild sufferers remaining undiagnosed should be taken into account when interpreting the results of this study.

Our findings suggest that meaningful approaches to the prevention and potential treatment of psoriasis and eczema (particularly psoriasis) might come from targeting adiposity levels in addition to the immune pathways in skin. This is supported by previous reports of weight loss improving psoriatic skin and joint disease[65-67]. There are various possible mechanisms linking obesity with skin inflammation due to functional changes within adipose tissue as well as quantitative effects, such as the increased production of inflammatory cytokines from adipose tissue[68]. Excess skin adipose tissue results in pro-inflammatory cytokine and hormone secretion. Cytokines such as TNFα and IL-6 are directly implicated in the pathology of psoriasis and are targets for some current highly effective treatments[69,70]. Leptin can increase keratinocyte proliferation and pro-inflammatory protein secretion which are characteristics of psoriasis[71], whilst the secretion of adiponectin, which is putatively anti-inflammatory[72] is reduced in the obese state. The skin of obese individuals shows features of impaired barrier function[73] which may predispose to atopic inflammation whilst impairment in lymphatic function may delay the clearance of inflammatory mediators[68]. Other mechanisms remain possible; however these are weakly researched. Our results, supporting a causal relationship, suggest this area warrants further detailed work.

In conclusion, our findings suggest a causal effect of BMI upon common inflammatory skin disease, which may carry important health implications. These results provide further evidence supporting the crucial importance of the management of obesity in patients with inflammatory skin diseases - more notably for psoriasis. Whereas this concept is already contained in relevant clinical guidelines, our data should provide even greater emphasis given the potential to yield substantial clinical benefits.

Our findings could potentially help inform the management or prevention of disease, and highlight how interventions for reducing obesity may reduce the burden of inflammatory skin disease; providing further rationale for trials of weight loss or weight loss agents in patients with psoriasis or eczema.

## Acknowledgements

This research has been conducted using data from the UK Biobank Resource (application number 10074) and the Nord-Trøndelag Health Study (the HUNT Study). The HUNT Study is a collaboration between HUNT Research Centre (Faculty of Medicine and Health Sciences, NTNU, Norwegian University of Science and Technology), Nord-Trøndelag County Council, Central Norway Regional Health Authority, and the Norwegian Institute of Public Health.

## Contributors

LP, BOA, TMF, SJB and NS conceived the study concept and managed the project. AB-A and JT performed statistical analysis with the UK Biobank Resource. BB and EHM performed statistical analysis with the HUNT dataset, whose genotypes were quality controlled and prepared primarily by LGF. SW conducted the literature review with support from AB-A, BB, SJB and LP. TP contributed towards the statistical analysis performed. ARW, SEJ and RB contributed towards the quality control of UK Biobank data. AB-A, SJB, NS, LP, and BB drafted the initial versions of the manuscript. All authors were involved in the interpretation of the data, and contributed to and approved the final version of the manuscript.

## Data sharing

The UK Biobank dataset used to conduct the research in this paper is available via application directly to the UK Biobank. Data from the HUNT Study used in research projects will when reasonably requested by others be made available upon request to the HUNT Data Access Committee. The HUNT data access information (available here: http://www.ntnu.edu/hunt/data) describes in detail the policy regarding data availability.

## REFERENCES

1 Nestle FO, Kaplan DH, Barker J. Psoriasis. N Engl J Med 2009;361:496–509. doi:10.1056/NEJMra0804595

2 Weidinger S, Novak N, Weidinger S, et al. Atopic dermatitis. Lancet 2016;387:1109–22. doi:10.1016/S0140-6736(15)00149-X

3 Armstrong AW, Harskamp CT, Armstrong EJ. The association between psoriasis and obesity: a systematic review and meta-analysis of observational studies. Nutr Diabetes 2012;2:e54. doi:10.1038/nutd.2012.26

4 Zhang A, Silverberg JI. Association of atopic dermatitis with being overweight and obese: a systematic review and metaanalysis. J Am Acad Dermatol 2015;72:606–16. doi:10.1016/j.jaad.2014.12.013

5 Jensen P, Zachariae C, Christensen R, et al. Effect of Weight Loss on the Severity of Psoriasis. JAMA Dermatology 2013;149:795. doi:10.1001/jamadermatol.2013.722

6 Egeberg A, Sørensen JA, Gislason GH, et al. Incidence and Prognosis of Psoriasis and Psoriatic Arthritis in Patients Undergoing Bariatric Surgery. JAMA Surg 2017;152:344. doi:10.1001/jamasurg.2016.4610

7 Al-Mutairi N, Nour T. The effect of weight reduction on treatment outcomes in obese patients with psoriasis on biologic therapy: a randomized controlled prospective trial. Expert Opin Biol Ther 2014;14:749–56. doi:10.1517/14712598.2014.900541

8 Lawlor DA, Harbord RM, Sterne JAC, et al. Mendelian randomization: using genes as instruments for making causal inferences in epidemiology. Stat Med 2008;27:1133–63. doi:10.1002/sim.3034

9 Burgess S, Butterworth A, Thompson SG. Mendelian randomization analysis with multiple genetic variants using summarized data. Genet Epidemiol 2013;37:658–65. doi:10.1002/gepi.21758

10 Smith GD, Ebrahim S. 'Mendelian randomization': Can genetic epidemiology contribute to understanding environmental determinants of disease? Int J Epidemiol 2003;32:1–22. doi:10.1093/ije/dyg070

11 Locke AE, Kahali B, Berndt SI, et al. Genetic studies of body mass index yield new insights for obesity biology. Nature 2015;518:197–206. doi:10.1038/nature14177

12 Tsoi LC, Stuart PE, Tian C, et al. Large scale meta-analysis characterizes genetic architecture for common psoriasis associated variants. Nat Commun 2017;8:15382. doi:10.1038/ncomms15382

13 Paternoster L, Standl M, Waage J, et al. Multi-ancestry genome-wide association study of 21,000 cases and 95,000 controls identifies new risk loci for atopic dermatitis. Nat Genet 2015;47:1449–56. doi:10.1038/ng.3424

14 Perry JRB, Weedon MN, Langenberg C, et al. Genetic evidence that raised sex hormone binding globulin (SHBG) levels reduce the risk of type 2 diabetes. Hum Mol Genet 2010;19:535–44. doi:10.1093/hmg/ddp522

15 Sudlow C, Gallacher J, Allen N, et al. UK Biobank: An Open Access Resource for Identifying the Causes of a Wide Range of Complex Diseases of Middle and Old Age. PLoS Med 2015;12:e1001779. doi:10.1371/journal.pmed.1001779

16 Krokstad S, Langhammer A, Hveem K, et al. Cohort profile: The HUNT study, Norway. Int J Epidemiol 2013;42:968–77. doi:10.1093/ije/dys095

17 Modalsli EH, Snekvik I, Åsvold BO, et al. Validity of self-reported psoriasis in a general population: The HUNT study, Norway. J. Invest. Dermatol. 2016;136:323–5. doi:10.1038/JID.2015.386

18 Wain L V, Shrine N, Miller S, et al. Novel insights into the genetics of smoking behaviour, lung function, and chronic obstructive pulmonary disease (UK BiLEVE): A genetic association study in UK Biobank. Lancet Respir Med 2015;3:769–81. doi:10.1016/S2213-2600(15)00283-0

19 Bycroft C, Freeman C, Petkova D, et al. Genome-wide genetic data on ~500,000 UK Biobank participants. doi.org 2017;:166298. doi:10.1101/166298

20 Mitchell R, Hemani G, Dudding T, et al. UK Biobank Genetic Data: MRC-IEU Quality Control, Version 1. doi.org 2017. doi:10.5523/bris.3074krb6t2frj29yh2b03x3wxj

21 Frayling TM, Beaumont R, Jones SE, et al. A common allele in FGF21 associated with preference for sugar consumption lowers body fat in the lower body and increases blood pressure. bioRxiv 2017;:214700. doi:10.1101/214700

22 Nielsen JB, Thorolfsdottir RB, Fritsche LG, et al. Genome-wide association study of 1 million people identifies 111 loci for atrial fibrillation. doi:http://dx.doi.org/10.1101/242149

23 Burgess S. Identifying the odds ratio estimated by a two-stage instrumental variable analysis with a logistic regression model. Stat Med 2013;32:4726–47. doi:10.1002/sim.5871

24 Burgess S, Small DS, Thompson SG. A review of instrumental variable estimators for Mendelian randomization. Stat Methods Med Res 2015;:962280215597579. doi:10.1177/0962280215597579

25 Burgess S, Scott RA, Timpson NJ, et al. Using published data in Mendelian randomization: A blueprint for efficient identification of causal risk factors. Eur J Epidemiol 2015; 30:543–52. doi:10.1007/s10654-015-0011-z

26 Yavorska OO, Burgess S. MendelianRandomization: an R package for performing Mendelian randomization analyses using summarized data. Int J Epidemiol Published Online First: 7 April 2017. doi:10.1093/ije/dyx034

27 Bowden J, Davey Smith G, Haycock PC, et al. Consistent Estimation in Mendelian Randomization with Some Invalid Instruments Using a Weighted Median Estimator. Genet Epidemiol 2016; 40:304–14. doi:10.1002/gepi.21965

28 Hartwig FP, Davey Smith G, Bowden J. Robust inference in summary data Mendelian randomization via the zero modal pleiotropy assumption. Int J Epidemiol 2017; 46:1985–98. doi:10.1093/ije/dyx102

29 Bowden J, Smith GD, Burgess S. Mendelian randomization with invalid instruments: Effect estimation and bias detection through Egger regression. Int J Epidemiol 2015; 44:512–25. doi:10.1093/ije/dyv080

30 Speliotes EK, Willer CJ, Berndt SI, et al. Association analyses of 249,796 individuals reveal 18 new loci associated with body mass index. Nat Genet 2010; 42:937–48. doi:10.1038/ng.686

31 Baiocchi M, Cheng J, Small DS. Instrumental variable methods for causal inference. Stat Med 2014; 33:2297–340. doi:10.1002/sim.6128

32 Duffy DL, Spelman LS, Martin NG. Psoriasis in Australian twins. J Am Acad Dermatol 1993; 29:428–34. doi:10.1016/0190-9622(93)70206-9

33 Naldi L, Chatenoud L, Linder D, et al. Cigarette smoking, body mass index, and stressful life events as risk factors for psoriasis: Results from an Italian case-control study. J Invest Dermatol 2005; 125:61–7. doi:10.1111/j.0022-202X.2005.23681.x

34 Neimann AL, Shin DB, Wang X, et al. Prevalence of cardiovascular risk factors in patients with psoriasis. J Am Acad Dermatol 2006; 55:829–35. doi:10.1016/j.jaad.2006.08.040

35 Cohen AD, Sherf M, Vidavsky L, et al. Association between psoriasis and the metabolic syndrome. A cross-sectional study. Dermatology 2008; 216:152–5. doi:10.1159/000111512

36 Setty AR, Curhan G, Choi HK. Obesity, waist circumference, weight change, and the risk of psoriasis in women: Nurses' Health Study II. ArchInternMed 2007; 167:1670–5. doi:10.1001/archinte.167.15.1670

37 Boccardi D, Menni S, Vecchia C La, et al. Overweight and childhood psoriasis. Br. J. Dermatol. 2009; 161:484–6. doi:10.1111/j.1365-2133.2009.09276.x

38 Driessen RJB, Boezeman JB, Van De Kerkhof PCM, et al. Cardiovascular risk factors in high-need psoriasis patients and its implications for biological therapies. J Dermatolog Treat 2009; 20:42–7. doi:10.1080/09546630802225702

39 Naldi L, Chatenoud L, Belloni A, et al. Medical history, drug exposure and the risk of psoriasis. Evidence from an Italian case-control study. Dermatology 2008; 216:125–30-2. doi:10.1159/000111509

40 Al-Mutairi N, Al-Farag S, Al-Mutairi A, et al. Comorbidities associated with psoriasis: An experience from the Middle East. J Dermatol 2010; 37:146–55. doi:10.1111/j.1346-8138.2009.00777.x

41 Schmitt J, Ford DE. Psoriasis is independently associated with psychiatric morbidity and adverse cardiovascular risk factors, but not with cardiovascular events in a population-based sample. J Eur Acad Dermatology Venereol 2010; 24:885–92. doi:10.1111/j.14683083.2009.03537.x

42 Koebnick C, Black MH, Smith N, et al. The Association of Psoriasis and Elevated Blood Lipids in Overweight and Obese Children. J Pediatr 2011;159:577–83. doi:10.1016/j.jpeds.2011.03.006

43 Zhu KJ, He SM, Zhang C, et al. Relationship of the body mass index and childhood psoriasis in a Chinese Han population: a hospital-based study. J Dermatol 2012;39:181–3. doi:10.1111/j.1346-8138.2011.01281.x

44 Kumar S, Han J, Li T, et al. Obesity, waist circumference, weight change and the risk of psoriasis in US women. J Eur Acad Dermatology Venereol 2013;27:1293–8. doi:10.1111/jdv.12001

45 Tseng HW, Lin HS, Lam HC. Co-morbidities in psoriasis: A hospital-based case-control study. J Eur Acad Dermatology Venereol 2013;27:1417–25. doi:10.1111/jdv.12028

46 Casagrande SS, Menke A, Cowie CC. No association between psoriasis and diabetes in the U.S. population. Diabetes Res Clin Pract 2014;104:e58–60. doi:10.1016/j.diabres.2014.04.009

47 Harpsøe MC, Basit S, Andersson M, et al. Body mass index and risk of autoimmune diseases: A study within the Danish National Birth Cohort. Int J Epidemiol 2014;43:843–55. doi:10.1093/ije/dyu045

48 Menegon DB, Pereira AG, Camerin AC, et al. Psoriasis and comorbidities in a southern Brazilian population: A case-control study. Int J Dermatol 2014;53:e518–25. doi:10.1111/ijd.12186

49 Votrubova J, Juzlova K, Smerhovsky Z, et al. Risk factors for comorbidities in Czech psoriatic patients: Results of a hospital-based case-control study. Biomed Pap 2014;158:288–94. doi:10.5507/bp.2013.062

50 Jacobi A, Langenbruch A, Purwins S, et al. Prevalence of Obesity in Patients with Psoriasis: Results of the National Study PsoHealth3. Dermatology 2015;231:231–8. doi:10.1159/000433528

51 Wolk K, Mallbris L, Larsson P, et al. Excessive body weight and smoking associates with a high risk of onset of plaque psoriasis. Acta Derm Venereol 2009;89:492–7. doi:10.2340/00015555-0711

52 Wolkenstein P, Revuz J, Roujeau JC, et al. Psoriasis in France and associated risk factors: Results of a case-control study based on a large community survey. Dermatology 2009;218:103–9. doi:10.1159/000182258

53 Ellison JA, Patel L, Kecojevic T, et al. Pattern of growth and adiposity from infancy to adulthood in atopic dermatitis. Br J Dermatol 2006;155:532–8. doi:10.1111/j.1365-2133.2006.07400.x

54 Kusunoki T, Morimoto T, Nishikomori R, et al. Obesity and the prevalence of allergic diseases in schoolchildren. Pediatr Allergy Immunol 2008;19:527–34. doi:10.1111/j.1399-3038.2007.00686.x

55 Machura E, Szczepanska M, Ziora K, et al. Evaluation of adipokines: apelin, visfatin, and resistin in children with atopic dermatitis. Mediators Inflamm 2013;2013:760691. doi:10.1155/2013/760691

56 Kim S, Lee J-Y, Oh J-Y, et al. The Association between Atopic Dermatitis and Depressive Symptoms in Korean Adults: The Fifth Korea National Health and Nutrition Examination Survey, 2007-2012. Korean J Fam Med 2015;36:261. doi:10.4082/kjfm.2015.36.6.261

57 Lee JH, Han K Do, Jung HM, et al. Association Between Obesity, Abdominal Obesity, and Adiposity and the Prevalence of Atopic Dermatitis in Young Korean Adults: the Korea National Health and Nutrition Examination Survey 2008-2010. Allergy Asthma Immunol Res 2016;8:107–14. doi:10.4168/aair.2016.8.2.107

58 Cole TJ, Bellizzi MC, Flegal KM, et al. Establishing a standard definition for child overweight and obesity worldwide: international survey. BMJ 2000;320:1240–3. doi:10.1136/BMJ.320.7244.1240

59 Palmer CNA, Irvine AD, Terron-Kwiatkowski A, et al. Common loss-of-function variants of the epidermal barrier protein filaggrin are a major predisposing factor for atopic dermatitis. Nat Genet 2006;38:441–6. doi:10.1038/ng1767

60 Danielsen K, Olsen AO, Wilsgaard T, et al. Is the prevalence of psoriasis increasing? A 30-year follow-up of a population-based cohort. Br J Dermatol 2013;168:1303–10. doi:10.1111/bjd.12230

61 Deckers IAG, McLean S, Linssen S, et al. Investigating International Time Trends in the Incidence and Prevalence of Atopic Eczema 1990-2010: A Systematic Review of Epidemiological Studies. PLoS One 2012;7:e39803. doi:10.1371/journal.pone.0039803

62 Statistics Team, NHS Digital NHS. Statistics on obesity, physical activity and diet. England, 2017. 2017. https://www.gov.uk/government/uploads/system/uploads/attachment_data/file/613532/obes-phys-acti-diet-eng-2017-rep.pdf (accessed 18 Jan 2018).

63 DaVeiga SP. Epidemiology of atopic dermatitis: a review. Allergy Asthma Proc 2012;33:227–34. doi:10.2500/aap.2012.33.3569

64 Monnereau C, Vogelezang S, Kruithof CJ, et al. Associations of genetic risk scores based on adult adiposity pathways with childhood growth and adiposity measures. BMC Genet 2016;17:120. doi:10.1186/s12863-016-0425-y

65 Maglio C, Peltonen M, Rudin A, et al. Bariatric Surgery and the Incidence of Psoriasis and Psoriatic Arthritis in the Swedish Obese Subjects Study. Obesity 2017;25:2068–73. doi:10.1002/oby.21955

66 Di Minno MND, Peluso R, Iervolino S, et al. Weight loss and achievement of minimal disease activity in patients with psoriatic arthritis starting treatment with tumour necrosis factor *α* blockers. Ann Rheum Dis 2014;73:1157–62. doi:10.1136/annrheumdis-2012-202812

67 Upala S, Sanguankeo A. Effect of lifestyle weight loss intervention on disease severity in patients with psoriasis: a systematic review and meta-analysis. Int J Obes 2015;39:1197–202. doi:10.1038/ijo.2015.64

68 Nakamizo S, Honda T, Kabashima K. Obesity and inflammatory skin diseases. Trends Immunother 2017;1:67–74. doi:10.24294/ti.v1i2.98

69 Dowlatshahi EA, van der Voort EA., Arends LR, et al. Markers of systemic inflammation in psoriasis: a systematic review and meta-analysis. Br J Dermatol 2013;169:266–82. doi:10.1111/bjd.12355

70 Sbidian E, Chaimani A, Garcia-Doval I, et al. Systemic pharmacological treatments for chronic plaque psoriasis: a network meta-analysis. Cochrane Database Syst Rev 2017;12:CD011535. doi:10.1002/14651858.CD011535.pub2

71 Stjernholm T, Ommen P, Langkilde A, et al. Leptin deficiency in mice counteracts imiquimod (IMQ)-induced psoriasis-like skin inflammation while leptin stimulation induces inflammation in human keratinocytes. Exp Dermatol 2017;26:338–45. doi:10.1111/exd.13149

72 Davidovici BB, Sattar N, Jörg PC, et al. Psoriasis and Systemic Inflammatory Diseases: Potential Mechanistic Links between Skin Disease and Co-Morbid Conditions. J Invest Dermatol 2010;130:1785–96. doi:10.1038/JID.2010.103

73 Löffler H, Aramaki JUN, Effendy I. The influence of body mass index on skin susceptibility to sodium lauryl sulphate. Ski Res Technol 2002;8:19–22. doi:10.1046/j.0909-752x

